# Pyronaridine and artesunate are potential antiviral drugs against COVID-19 and influenza

**DOI:** 10.1101/2020.07.28.225102

**Authors:** Joon-Yong Bae, Gee Eun Lee, Heedo Park, Juyoung Cho, Yung-Eui Kim, Joo-Yeon Lee, Chung Ju, Won-Ki Kim, Jin Il Kim, Man-Seong Park

## Abstract

Since the first human case was reported in Wuhan Province, China in December 2019, SARS-CoV-2 has caused millions of human infections in more than 200 countries worldwide with an approximately 4.01% case-fatality rate (as of 27 July, 2020; based on a WHO situation report), and COVID-19 pandemic has paralyzed our global community. Even though a few candidate drugs, such as remdesivir (a broad antiviral prodrug) and hydroxychloroquine, have been investigated in human clinical trials, their therapeutic efficacy needs to be clarified further to be used to treat COVID-19 patients. Here we show that pyronaridine and artesunate, which are the chemical components of anti-malarial drug Pyramax^®^, exhibit antiviral activity against SARS-CoV-2 and influenza viruses. In human lung epithelial (Calu-3) cells, pyronaridine and artesunate were highly effective against SARS-CoV-2 while hydroxychloroquine did not show any effect at concentrations of less than 100 μM. In viral growth kinetics, both pyronaridine and artesunate inhibited the growth of SARS-CoV-2 and seasonal influenza A virus in Calu-3 cells. Taken together, we suggest that artesunate and pyronaridine might be effective drug candidates for use in human patients with COVID-19 and/or influenza, which may co-circulate during this coming winter season.

## Main Text

Severe acute respiratory syndrome coronavirus 2 (SARS-CoV-2) has caused a pandemic, and more than 16 million cases of coronavirus disease 2019 (COVID-19) have been reported across at least 200 countries and territories worldwide, resulting in approximately 646,000 deaths (as of 27 July 2020) (1). To treat COVID-19 patients and to prepare for a possible second wave in advance, effective antiviral drugs against SARS-CoV-2 are urgently needed.

The antimalarial drugs chloroquine and hydroxychloroquine have been testified against infections caused by many DNA and RNA viruses, including human coronaviruses (2). However, the results of clinical studies of those drugs for the treatment of COVID-19 have not been favorable (3). Here, we are the first to report the potential of another antimalarial drug, Pyramax^®^, a fixed-dose combination of pyronaridine and artesunate that is currently under a phase II clinical trial in Republic of Korea (4), for COVID-19 treatment. The combination of pyronaridine and artesunate has previously been reported to exhibit broad-spectrum antiviral activity (5, 6). In particular, pyronaridine showed more effective protection and a greater reduction in viral load than other lysosomotropic antimalarial drugs in Ebola virus-challenged mice and guinea pigs (7, 8). Additionally, artesunate reduced the viral load by 20-fold in cytomegalovirus-infected rats (9). In this study, we show that pyronaridine and artesunate also exhibit antiviral activity against SARS-CoV-2 and influenza viruses.

Antiviral effects of pyronaridine and artesunate were evaluated by viral RNA loads, cell viability, viral growth kinetics, and time-of-drug-addition assays. First, in Vero cells, pyronaridine inhibited SARS-CoV-2 replication with a half-maximal inhibitory concentration (IC_50_) of 1.084 μM, a half-maximal cytotoxic concentration (CC_50_) of 37.09 μM and a selectivity index (SI) of 34.22 at 24 hours post infection (hpi). The corresponding values for artesunate were an IC_50_ of 53.06 μM, a CC_50_ of higher than 100 μM (> 100 μM) and an SI of > 1.885. However, the inhibitory effects of pyronaridine and artesunate against SARS-CoV-2 in Vero cells were not vastly superior to those of hydroxychloroquine, given the results determined at 24 and 48 hpi (Fig. 1A). Interestingly, in Calu-3 cells, while hydroxychloroquine did not show antiviral effect against SARS-CoV-2 at concentrations of less than 100 μM, pyronaridine and artesunate were highly effective against SARS-CoV-2. The inhibitory effects of artesunate in Calu-3 cells (at 24 hpi: IC_50_ 1.76 μM, CC_50_ > 100 μM, and SI > 56.82; and 48 hpi: IC_50_ 0.453 μM, CC_50_ > 100 μM, and SI > 220.8) were notably better than those of artesunate in Vero cells and were also better than those of pyronaridine in Calu-3 cells (at 24 hpi: IC_50_ 6.413 μM, CC_50_ 43.08 μM, and SI 6.718; and at 48 hpi: IC_50_ 8.577 μM, CC_50_ > 100 μM, and SI > 11.66) (Fig. 1B). By determining viral growth kinetics in Calu-3 cells, we confirmed these two drugs could reduce viral replication in a dose-dependent manner whereas hydroxychloroquine showed no antiviral effects at 1.56-50 mM concentrations (Fig. 1C), and the results of the time-of-addition assay might indicate that artesunate functions at a stage after viral entry (Fig. 1D). As demonstrated in the monotherapy approaches (Fig. 1A-C), combinations of various concentrations of pyronaridine and artesunate also exhibited antiviral efficacy against SARS-CoV-2, and of these drugs, artesunate demonstrated much better effects against the growth of seasonal influenza A(H1N1) virus in Calu-3 cells (Fig. 1E), which highlights the possible broad-spectrum use of artesunate against COVID-19 and influenza.

**Fig. 1.**
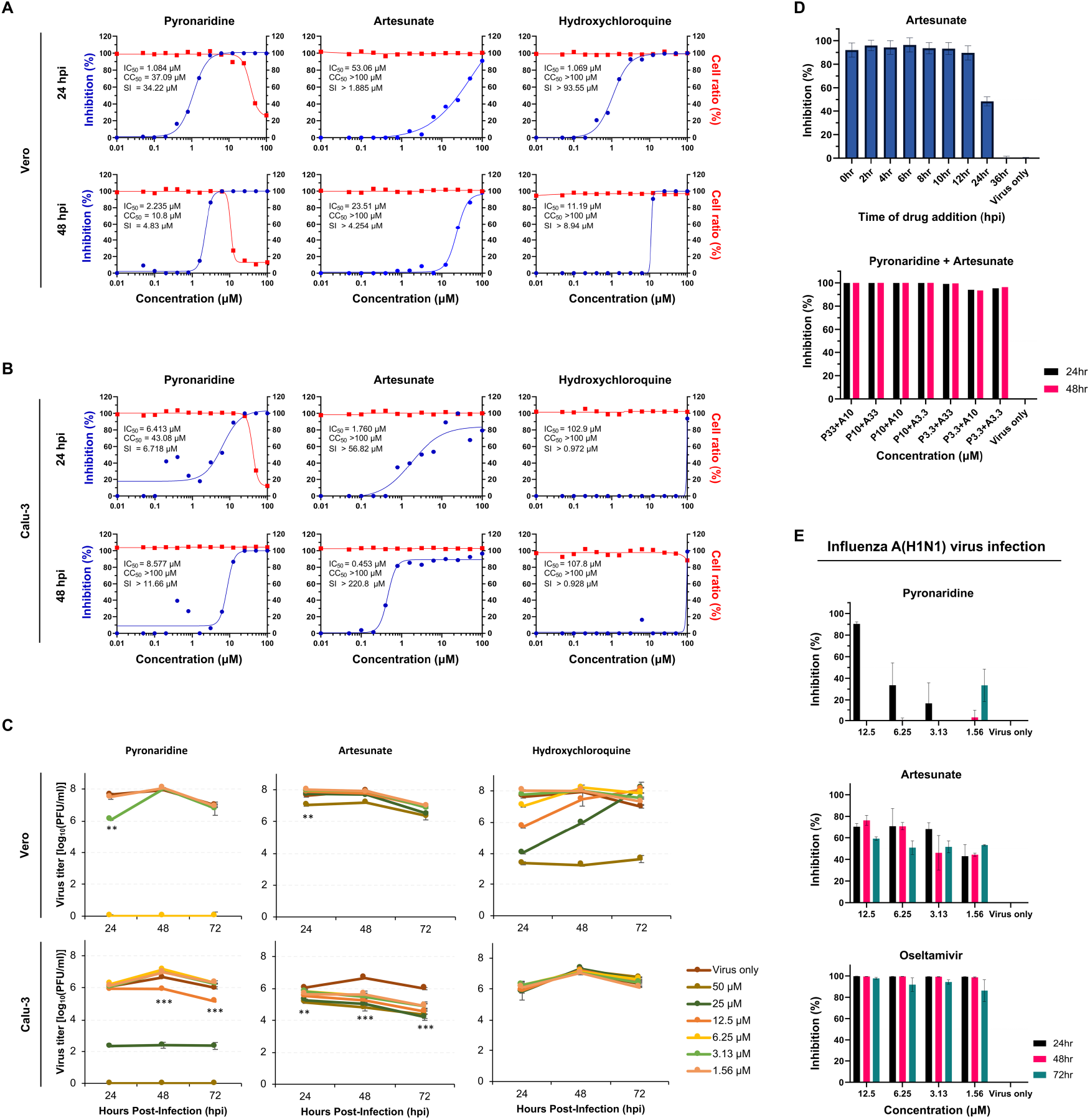
Antiviral efficacy of the pyronaridine and artesunate against SARS-CoV-2 and influenza viruses in cells. (A, B) The antiviral effects of pyronaridine, artesunate, or hydroxychloroquine was evaluated by qRT-PCR using the cell supernatants infected with SARS-CoV-2. The cytotoxicity of these drugs was measured by a WST-1 cell proliferation assay using the same cell supernatants. (C) The antiviral effects of pyronaridine, artesunate, and hydroxychloroquine on SARS-CoV-2 replication were evaluated in Vero or Calu-3 cells. The virus-only group was treated with cell culture medium. (D) The antiviral effects of artesunate was measured by a time-to-addition assay (with artesunate intervention at different time points) in Calu-3 cells. The virus-only group was treated with cell culture medium. Combination effects of various concentrations of pyronaridine and artesunate on SARS-CoV-2 was evaluated by qRT-PCR using the Vero cell supernatants. The virus-only group was treated with cell culture medium. (E) The antiviral effects of pyronaridine and artesunate against human seasonal influenza A(H1N1) virus were evaluated in Calu-3 cells. Oseltamivir was used as a control drug. The virus-only group was treated with cell culture medium.

The antiviral effects of both artesunate and its active metabolite dihydroartemisinin (DHA) against DNA viruses have been well described and appear to be mediated through NF-kB and Sp1, which are involved in the type-1 interferon (IFN) pathway (7). In addition to having antiviral effects, artesunate has been reported to have direct protective effects against sepsis-induced lung injury by inhibiting the Toll-like receptor 4 (TLR4) inflammatory signaling pathway in mice (10). The mode of action described above is consistent with our data, which indicate that the antiviral efficacy of artesunate against SARS-CoV-2 in Calu-3 cells is far superior to that in Vero cells, which are well known for their type-1 IFN pathway deficiency. Notably, the inhibitory effects of artesunate against SARS-CoV-2 infection were worse than those of hydroxychloroquine in Vero cells but better than those of hydroxychloroquine in Calu-3 cells, which are derived from human airway epithelial cells and thus may be more representative of the susceptible cells in actual human airway infection than Vero cells.

Our data demonstrate a new possibility of repurposing antimalarial drugs against both SARS-CoV-2 and influenza A viruses. Fortunately, artesunate is an FDA-approved drug for malaria with no safety issues identified to date. Our data suggest that artesunate has the potential to be used as a primary selective drug if COVID-19 occurs during the upcoming influenza season in winter. In addition, an *in vivo* study needs to be conducted to evaluate the efficacy of artesunate against SARS-CoV-2 and influenza viruses. If its antiviral efficacy is proven effective in clinical trials, the combination of artesunate and pyronaridine could be an effective drug for use in human patients with COVID-19 and/or influenza, which may circulate mainly during the winter season.

## Materials and Methods

### Cells and viruses

Vero, Calu-3, and Madin-Darby canine kidney (MDCK) cells were purchased from the Korean Cell Line Bank (Seoul, Republic of Korea). The cells were maintained in Dulbecco’s modified Eagle’s medium (DMEM; Gibco, Thermo Fisher Scientific, Waltham, MA) supplemented with 10% fetal bovine serum (FBS; Serana, Pessin, Germany) and penicillinstreptomycin (Gibco). SARS-CoV-2 (hCoV-19/South Korea/KCDC03/2020) was provided by the Korea Centers for Disease Control and Prevention (Osong, Republic of Korea) and prepared by propagation in Vero cells after plaque purification. Influenza A(H1N1) virus (A/Korea/01/2009, NCBI:txid644289) was prepared by propagation in MDCK cells after plaque purification.

### Chemicals

Pyronaridine and artesunate were provided from Shinpoong Pharmaceutical Co., LTD (Seoul, Republic of Korea). Hydroxychloroquine sulfate was purchased from Sigma-Aldrich (St. Louis, MO). Oseltamivir carboxylate was purchased from Toronto Research Chemicals (Ontario, Canada). The chemicals were dissolved in dimethyl sulfoxide (DMSO; Sigma-Aldrich) or deionized water to achieve a final concentration of 10 mM.

### Real-time RT-PCR of SARS-CoV-2 RNAs

Vero or Calu-3 cells were infected with SARS-CoV-2 at 0.01 or 0.1 of multiplicity of infection (MOI), respectively. After 1 h, the cells were washed three times with DMEM and maintained with DMEM containing 2% FBS (2% FBS-DMEM).

Subsequently, the cells were treated with serial two-fold dilutions of pyronaridine, artesunate, or hydroxychloroquine (starting concentration = 100 μM). At 24 and 48 hpi, viral RNAs in the cell supernatants were quantified by real-time RT-PCR. Briefly, cell supernatants were harvested for RNA extraction with a Maxwell RSV Viral Total Nucleic Acid Purification Kit (Promega, Madison, WI). Then, a SARS-CoV-2 RdRp region was amplified using forward (5’-GTGARATGGTCATGTGTGGCGG) and reverse (5’-CARATGTTAAASACACTATTAGCATA) primers and a probe (5’-FAM-CAGGTGGAACCTCATCAGGAGATGC – BHQI) (11). 50% inhibitory concentration (IC_50_) of pyronaridine, artesunate, or hydroxychloroquine was calculated using GraphPad Prism 8 (GraphPad, San Diego, CA).

### Cell viability assay

A confluent monolayer of Vero or Calu-3 cells was maintained with 2% FBS-DMEM. Then, serial two-fold dilutions of pyronaridine, artesunate, or hydroxychloroquine (starting concentration = 100 μM) were added to each well. DMSO was used as a control. At 24 and 48 hpi, 10 μl of WST-1 was added to each well. After 2 h, the cell viability was measured, and 50% cytotoxic concentration (CC_50_) was calculated using GraphPad Prism 8 (GraphPad).

### Growth kinetics of SARS-CoV-2

Monolayered Vero or Calu-3 cells were inoculated with SARS-CoV-2 at a 0.01 or 0.1 MOI for 1 h, respectively. Then, the inoculum was discarded, and the cells were maintained with 2% FBS-DMEM. Serial two-fold dilution from 50 μM to 1.56 μM of chemicals were added to the cell supernatants, and the cell supernatants were harvested at 24, 48, and 72 hpi for virus titration by the plaque assay in Vero cells. A virus only group was treated with DMEM.

### Plaque assay

Monolayered Vero cells prepared in advance was inoculated with the virus-infected cell supernatants. After 1 h, the cells were overlaid with DMEM-F12 (Sigma-Aldrich) containing 2% agarose. After 72 h, the cells were stained with crystal violet (Georgia Chemicals Inc., Norcross, GA). For the H1N1 influenza virus, MDCK cells were used, instead of Vero cells.

### Combination therapy

Monolayered Vero cells was inoculated with SARS-CoV-2 at a 0.01 MOI. The inoculum was discarded after 1 h, and the cells were maintained with DMEM containing 2 % FBS. Various concentration combinations of pyronaridine and artesunate (for pyronaridine and artesunate concentrations: 33 μM + 10 μM; 10 μM + 33 μM; 10 μM + 10 μM; 10 μM + 3.3 μM; 3.3 μM + 33 μM; 3.3 μM + 10 μM; and 3.3 μM + 3.3 μM) were then added to the cell supernatants, and the cell supernatants harvested at 24 and 48 hpi were used for the qRT-PCR analysis.

### Time-to-addition assay

Monolayered Calu-3 cells was inoculated with SARS-CoV-2 at a 0.1 MOI. The inoculum was discarded after 1 h, and the cells were washed with DMEM three times and maintained with DMEM containing 2 % FBS. Artesunate (12.5 μM) were then added to the cell supernatants at 0, 2, 4, 6, 8, 10, 12, 24, and 36 hpi. The cell supernatants were harvested 48 h later from each drug-addition time point, and virus titers were determined by the plaque assay in Vero cells.

### Yield reduction of influenza virus

Monolayered Calu-3 cells was inoculated with influenza virus at a 0.1 MOI for 1 h. Then, the inoculum was discarded, and the cells were maintained with 0.3% FBS-DMEM. Serial two-fold dilution from 12.5 μM to 1.56 μM of pyronaridine, artesunate, or oseltamivir carboxylate were added to the cell supernatants, and the cell supernatants were harvested at 24, 48, and 72 hpi for virus titration by the plaque assay in MDCK cells. A virus only group was treated with DMEM.

The data, associated protocols, code, and materials described in the manuscript are available upon request.

## Acknowledgments

This study was supported by a grant from the National Research Foundation of Korea (NRF) funded by the Ministry of Science and ICT, Republic of Korea (Grant No. NRF-2017M3A9E4061995).

